# *SMART MRS*: A Simulated MEGA-PRESS Artifacts Toolbox for GABA-edited MRS

**DOI:** 10.1101/2024.09.19.612894

**Authors:** Hanna Bugler, Amirmohammad Shamaei, Roberto Souza, Ashley D. Harris

## Abstract

**Purpose:** To create a Python-based toolbox to simulate commonly occurring artifacts for single voxel Gamma-Aminobutyric Acid (GABA)-edited Magnetic Resonance Spectroscopy (MRS) data.

**Methods:** The toolbox was designed to maximize user flexibility and contains artifact, applied, input/output (I/O), and support functions. The artifact functions can produce spurious echoes, eddy currents, nuisance peaks, line broadening, baseline contamination, linear frequency drifts, and frequency and phase shift artifacts. Applied functions combine or apply specific parameter values to produce recognizable effects such as lipid peak and motion contamination. I/O and support functions provide additional functionality to accommodate different kinds of input data (MATLAB FID-A .mat files, nifti-mrs files), which vary by domain (time vs. frequency), collection type (edited vs. non-edited) and scale.

**Results:** Users may provide some or all of the required artifact function parameter values which will result in artifacts of different appearances. Visual assessment confirms the resemblance of simulated compared to *in vivo* produced artifacts.

**Conclusion:** Our pip installable Python artifact simulated toolbox *SMART_MRS* is meant to enhance the diversity and quality of existing simulated edited-MRS data and is complementary to existing MRS simulation software.

## 1. INTRODUCTION

Magnetic resonance spectroscopy (MRS) is a method to non-invasively measure metabolite concentrations *in vivo*. In a typical spectrum, Gamma Aminobutyric Acid is overlapped by more abundant metabolites hindering its quantification. To overcome this, GABA-editing can be used to remove the overlapping creatine peak [1,2]. In this approach, an editing pulse applied at 1.9 ppm modulates the coupled GABA signal at 3 ppm (edit-ON condition). This is interleaved with an edit-OFF condition with no editing pulse (or an editing pulse applied outside the spectral range of interest). The creatine peak is unaffected in both conditions so after subtracting the edit-OFF transients from the edit-ON transients, the creatine signal is removed leaving the GABA signal. Because there is a macromolecule signal coupled between 1.7 and 3 ppm, which is also co-edited by the editing pulse at 1.9 ppm, the difference signal includes macromolecules and is therefore often referred to as GABA+ [3].

GABA+-edited MRS data is at-risk for artifacts seen in conventional (non-edited MRS) as well as some specific artifacts due to the frequency selective editing pulses and the subtraction of the edit-ON – edit-OFF pairs [4]. Artifacts which may affect GABA-edited MRS data include spurious echoes, eddy currents, and lipid contamination which are further described below.

### Spurious Echoes

Gradient spoiling aims to mitigate the refocusing of signals from outside the voxel. When gradient spoiling is imperfect, typically in combination with B0 inhomogeneities, it can result in the refocusing of spurious or unwanted echoes also known as a ghost artifact [5,6,7]. The oscillations from the contaminated, unwanted echo cannot always be removed from the signal of interest, in which case the affected transient may need to be discarded [8,9]. Altering the amplitude or timing of the spoiling gradients [7,8] or using outer-volume suppression (OVS) [6,10] can reduce the prevalence of unwanted echoes at acquisition but spurious echoes can still occur.

### Eddy Currents

Gradients used for spatial localization and water suppression can induce current in conductive elements of the scanner causing short abrupt shifts in the B0 field called eddy currents [8,11,12]. The GABA+-edited frequency selective editing pulses are vulnerable to eddy currents, due to their narrow bandwidth, resulting in spectra with asymmetric altered line shapes [7,13]. The time dependency of the artifact allows for correction either by subtracting a phase function matching the time dependency of the artifact [4,8,12] or accounting for the artifact while fitting the spectrum during quantification [7]. While these methods can recover eddy current contaminated data, their performance can be limited in the presence of other artifacts [7].

### Lipid contamination

Nuisance peak signals can be heterogeneous in appearance (singlet, doublet, triplet, etc.) and causes challenges in signal modelling. In single voxel spectroscopy (SVS), lipid contamination can occur in cortical voxels due to the proximity of the skull and imperfect voxel placement or participant motion [7,9]. Lipid contamination can be corrected using convolution-difference filtering and singular value decomposition, as used to mitigate poor water suppression [9]. However, unlike water, lipid peaks tend to have broad line shapes and can overlap with metabolites of interest making their removal more challenging [8]. Therefore, lipids can be accounted for in the fitting, but these methods are imperfect. For GABA+-edited MRS, lipid contamination does not generally directly affect the 3 ppm peak but the lipid signal at 1.5 ppm in the baseline can have a negative impact on preprocessing performance and fit quality [8,14].

### Baseline changes

A poor baseline can be due to a wide variety of known factors such as poor localization, outer volume signal, insufficient water suppression, hardware imperfections and other artifacts [9]. However, broadly, baseline imperfections/artifacts are described as smooth [9] and can be fitted by a smooth spline function [10].

### Motion contamination

Single voxel spectroscopy motion can be difficult to detect [15]. Motion (either physiological or head movement) can shift the original voxel placement potentially leading to the presence of outer-volume signal (i.e. cranial lipids) and/or alter the frequency and phase of the acquired data [9]. However, frequency shifts are not unique to motion and can also arise from B0 drift, for example gradient induced frequency drift [16]. Due to the lengthy scan times required to achieve satisfactory SNR (approximately 320 transients taking 10 minutes) [3], GABA+-editing is at an increased risk of artifacts due to participant motion [14,17]. Edited-MRS is particularly sensitive to frequency (and phase) changes as they can cause subtraction artifacts [2]. Other possible secondary effects of motion can include incorrect peak shapes, nuisance signals and accompanied baseline changes, line broadening, reduced peak areas from discordant peak phases, reduced water suppression and ultimately incorrect quantification [9,15]. Frequency and phase correction can re-align some degree of drift between transients (either from a small degree of motion or gradients induced frequency drift) [18]. However, large frequency and phase shifts can impact the editing of the GABA signal in addition to causing large subtraction artifacts [19].

Research to improve the performance of preprocessing methods aims to eliminate or reduce the effect of artifacts on metabolite quantification. Simulations provide a mechanism to test the performance of new preprocessing methods as they can include a known target. Furthermore, open access to in vivo data with artifacts is limited because poor-quality data is traditionally discarded. While simulation tools do exist [4,20], they do not (i) consider the specific needs of editing, and (ii) do not provide specific methods for simulating motion.

In this work, we propose an open-source Python-based artifact simulation toolbox for GABA+-edited MRS data. Though many artifacts exist, the following commonly occurring artifacts were selected to be included in the toolbox due to their prominence in and impact on GABA+-edited MRS: eddy currents, spurious echoes, line broadening, baseline contamination, frequency drift, frequency and phase shifts, lipid (nuisance) peaks, various degrees of motion. This toolbox is available on GitHub for widespread use and available as a pip installable Python package: SMART_MRS.

## 2. MATERIALS AND METHODS

### 2.1 Toolbox Overview

The *SMART MRS* (Simulated MEGA-PRESS ARTifacts for MRS) toolbox for GABA+-edited MRS is a set of Python-based functions that provide easy integration of commonly occurring artifacts into previously simulated artifact free GABA-edited MRS data as seen in Figure 1. Users are expected to provide one or many artifact-free simulated GABA-edited MRS FIDs, at least one of each ‘ON’ and ‘OFF’ subspectrum, which can be imported to Python with our toolbox I/O functions or manually loaded with the format [number of transients, number of spectral points]. Users are then recommended to apply functions from the Support module to interleave their subspectra, to more accurately represent the order of an acquired scan and scale their data to increase the applicability of Artifact function default parameter values. With the data prepared, users can then:

**Figure 1.**
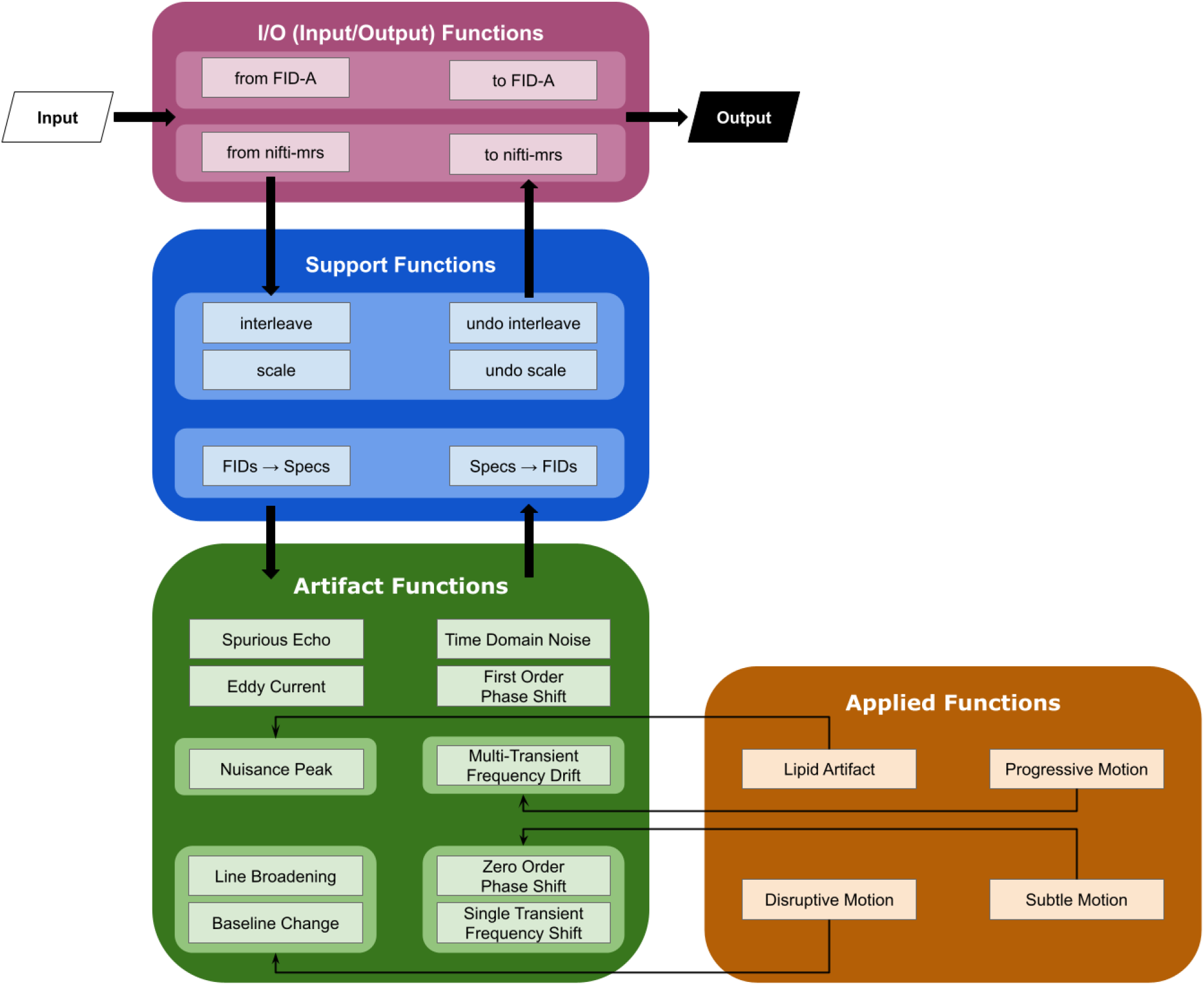
SMART MRS toolbox workflow divided into functional categories. I/O functions support the import and export of data into Python for use of the toolbox. Support functions enable effective use of the artifact and applied functions of the toolbox. Artifact functions add specific artifacts to existing simulated data. Applied functions provide specific use cases of artifact functions to mimic more complex events such as differing degrees of motion.

- Apply an Artifact function with user-selected parameter values
- Apply an Artifact function with randomized default parameter within a pre-determined range
- Apply an Artifact function with a mix of user-selected parameter values and randomized default parameter within a pre-determined range
- Apply an Applied function which is one or more Artifact functions with pre-determined default parameter values

Users can repeat this process sequentially adding one or many artifacts to their simulated data. When finished, users can apply functions from the I/O module to save their artifact contaminated data in their original imported format.

Functions within the *SMART MRS* toolbox are separated into four functional categories:

1. <u>Artifact Functions</u> are used to insert an artifact. Each function defines an artifact; to insert multiple different artifacts, these functions can be called serially. Currently defined artifacts include spurious echoes, eddy currents, nuisance peaks, line broadening, baseline contamination, linear frequency drifts, and frequency and phase shift. Default parameter values are provided for each artifact function and the user can change parameters to set the scale and appearance of the artifact as desired. All artifact functions include four general parameters for functionality. All artifact functions come with the following functional parameters in addition to artifact specific parameters:
  a. <u>Cluster:</u> Boolean value assigned to dictate whether the transients containing artifacts will be sequential within a scan (default is false).
  b. <u>Locs:</u> list of integer values corresponding to the transient numbers for the artifact containing transients (default is none with transient number(s) selected randomly at run-time).
  c. <u>Nmb:</u> integer value corresponding to the number of artifacts (default value is 1). In the case where Locs and Nmb do not agree, Nmb is prioritized over Locs and the latter is re-allocated at run-time.
  d. <u>Echo:</u> Boolean value assigned to dictate whether to print which default function parameters were used compared to those which were assigned by the user (default is False).
2. <u>Applied Functions</u> address artifacts that (a) rely on specific parameters such as macromolecule or lipid peaks appearing at a specific frequency or (b) are more complex because these artifacts have multiple effects on the spectrum such as motion that results in line broadening and frequency and phase offsets.
3. <u>Input/Output (I/O) Functions</u> are used for importing artifact-free simulated data from popular formats such as FID-A [4] .mat structures and nifti-mrs file format [21] to Python and exporting artifact contaminated simulated data from Python to these same formats. These formats were also selected for their flexibility to use data from other sources such as JMRUI (which can be opened with FID-A [4]) [22], OSPREY (compatible with nifti-mrs) [23], FSL-MRS (compatible with nifti-mrs) [24], and vendor specific data (which can be saved as a nifti-mrs file with spec2nii library [25]).
4. <u>Support Functions</u> are used for supporting the integration of artifacts into the simulated data but are not inserting artifacts themselves. Examples include domain conversion to convert between time and frequency domains, interleaving to insert artifacts on specified subspectra, and scaling functions for data not scaled within a standard range.

A list of all functions and their documentation can be found in the supplementary materials.

### 2.2 Artifact Functions

#### 2.2.1 Spurious Echo Artifact Simulation

In the *SMART MRS* toolbox, spurious echo artifacts are modelled in the time domain using *add_spur_echo_artifact()* by the following equation (Eq.1) adapted from Berrington et al [6]:

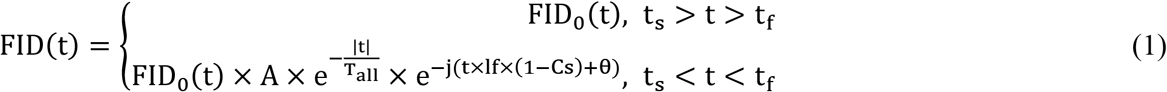

where *t* is time in the *FID, t*_*s*_ and *t*_*f*_ define the start and end point of the spurious echo, *FID*_*0*_(t) is a vector of the time domain signal as function of time, *A* is the amplitude of the artifact in the time domain, *T_all* is the length of the *FID, lf* is the Larmor frequency of hydrogen in Hz, *Cs* is the center of the echo artifact in the frequency domain in ppm, and *θ* is the assigned phase of the spurious echo (radians).

#### 2.2.2 Eddy Current Artifact Simulation

Eddy current artifacts are modelled in the time domain using *add_eddy_current_artifact()* by the following equation (Eq.2) adapted from FID-A [4]:

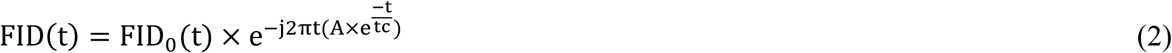

where *t* is time in the *FID, FID*_*0*_(t) is a vector of the time domain signal as function of time, *A* is the amplitude of the artifact and *t*_*c*_ is the time constant of the eddy current decay artifact in milliseconds.

#### 2.2.3 Nuisance Peak Artifact Simulation

The nuisance peak is modelled in the time domain by one of the following three equations found in *add_nuisance_peak()* based on user selection: Gaussian peak shape (Eq.3), Lorentzian peak shape (Eq.4) or pseudo-Voigt peak shape (Eq.5)

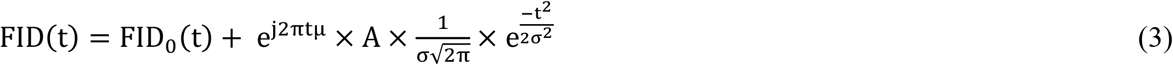

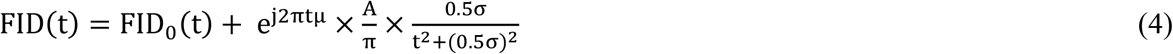

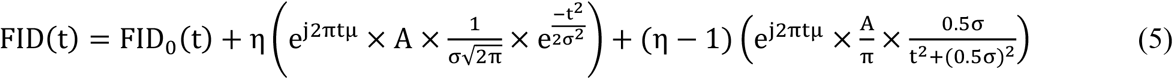

where *t* is time in the *FID, FID*_*0*_(t) is a vector of the time domain signal as function of time, *A* denotes the amplitude of the nuisance peak, σ is the FWHM of the nuisance peak, μ denotes the center frequency of the nuisance peak, and η is the Gaussian peak fraction used in a pseudo-Voigt peak. Users create profiles to model different peak shapes by creating a Python dictionary containing the desired peak type (“G” for Gaussian (Eq.3), “L” for Lorentzian (Eq.4), and “V” for pseudo-Voigt (Eq.5)), the amplitude, width (FWHM in ppm) and resonant frequency of each peak, and whether the transients belong to an edited sequence.

#### 2.2.4 Baseline Artifact Simulation

This toolbox contains functions within *add_baseline()* to add a baseline signal contamination in the frequency domain as modelled by one of the following two user-selected equations: sine wave (Eq. 6) or sinc wave (Eq. 7):

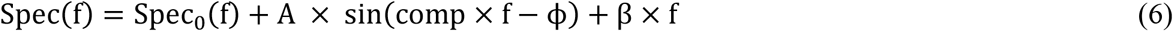

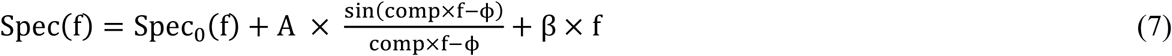

where *f* contains the frequency axis values of the *Spec, Spec*_*0*_(f) is a vector of the frequency domain signal as a function of frequency, *A* denotes the amplitude of each baseline component, *comp* denotes the frequency of the baseline cycles, φ denotes the phase of the baseline, and β denotes the slope of the baseline. As with the nuisance artifacts, users can create profiles to model different baselines by creating a Python dictionary containing the desired base type. Here, the user can also specify a number of wave functions to be included in the generation of the final baseline artifact, which may be either summed or summed and spline fit.

#### 2.2.5 Line Broadening Artifact Simulation

The toolbox contains a function *add_linebroad()* to broaden the lineshape of the existing signal, modelled in the time domain by the following equation (Eq.8):

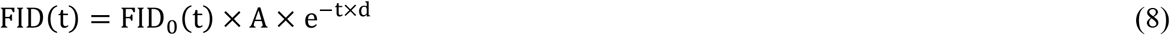

where *t* contains the time values of the *FID, FID*_*0*_(t) is a vector of the time domain signal as function of time, *A* denotes the amplitude of the decaying function, and *d* is the dampening coefficient and controls the degree of smoothing.

#### 2.2.6 Linear Frequency Drift

The toolbox contains a function *add_freq_drift_linear()* to add a linear frequency drift which spans multiple transients. This is modelled in the time domain by the following equation (Eq.9):

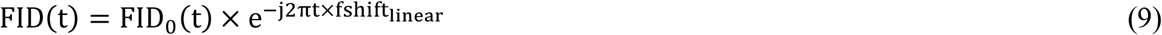

where *t* is the time of the *FID, FID*_*0*_(t) is a vector of the time domain signal as function of time, *f*_*shift_linear*_ is a set of values to define a linear shift across a set of transients. Within this function, a total frequency drift is defined as the total frequency change across a defined number of transients. Additionally, the magnitude of deviations from this linear function is also defined. The default values consist of a random linear drift value as a function of the number of transients between ±0.075×number of transients [19] and a deviation of 0.001 Hz.

#### 2.2.6 Frequency and Phase Shifts

The toolbox contains functions model in the time domain frequency and phase shifts divided into the following: frequency shifts using *add_freq_shift()* (Eq.10), zero order phase shifts using *add_zero_order_phase_shift()* (Eq.11), and first order phase shifts using *add_first_order_phase_shift()* (Eq.12) [4]:

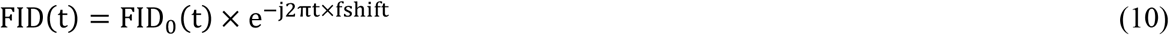

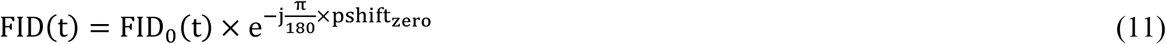

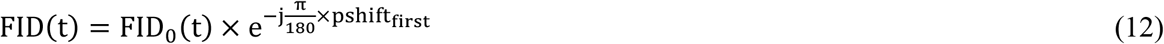

where *t* is time in the *FID, FID*_*0*_(t) is a vector of the time domain signal as function of time, *fshift* and *pshift*_*zero*_ are a set of normally distributed values with zero mean and standard deviation. The first order phase equation’s *pshift*_*first*_ is a function of frequency and a user-selected time shift (in ms). For *fshift* and *pshift*_*zero*_, the user may provide values for the standard deviation of the normal distribution from which frequency and phase shifts are sampled from. *fshift* affects the magnitude and direction of the frequency translation and *pshift*_*zero*_ affects the rotation of the peaks along the real-complex plane displayed as a change in peak shape and/or amplitude.

#### 2.2.8 Complex Time Domain Noise

The toolbox contains a function *add_time_domain_noise()* to produce complex time domain noise implemented as independent normal noise distributions applied to each the real and imaginary portion of the signal (Eq.13):

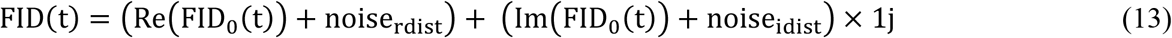

where *t* is time in the *FID, FID*_*0*_(t) is a vector of the time domain signal as function of time, Re() is the real portion of the signal, Im() is the imaginary portion of the signal, noise_rdist_ is the normally distributed noise applied to the real portion of the signal and noise_idist_ is the normally distributed noise applied to the imaginary portion of the signal.

### 2.3 Applied Functions

#### 2.3.1 Lipid Peak Artifact Simulation

In the *SMART MRS* toolbox, lipid artifacts are modelled by applying a lipid peak profile to the nuisance peak function. The peak profile is preset to be a singlet Gaussian peak with amplitude 0.025, width of 0.2 ppm, and resonant frequency of 1.5 ppm to lipid contamination often seen in GABA-edited MRS [14].

#### 2.3.2 Motion Artifact Simulation

There are various types of motion that will impact spectral quality to different degrees and are addressed in different ways. We have differentiated three types of motion that are mimicked using combinations of the aforementioned artifacts:

1. <u>‘Subtle’</u>: small discrete motions (*e*.*g*., jittering/fidgeting) that may be throughout the scan or during a defined period which are modelled by the *add_freq_shift()* and *add_zero_order_phase_shift()* functions.
2. <u>‘Progressive’</u>: systematic motion throughout the scan (e.g., head drifting as a subject falls asleep) modelled by the *add_freq_drift_linear()* function.
3. <u>‘Disruptive’</u>: large discrete motion (e.g., cough) affecting only a few transients modelled by the *add_linebroad()* and *add_baseline()* functions.

### 2.4 Support Functions

#### 2.4.1 Import Functions

The import/export functions facilitate the use of this toolbox for data simulated using other software such as FID-A [4]. Functions have been developed to import and export data with a FID-A structure and a nifti-mrs structure.

#### 2.4.2 Domain Conversion Functions

With all artifact functions requiring the time domain data, users are expected to keep data in the time domain. However, visualization in the frequency domain may be desirable so a pair of functions to convert between the frequency and time domain has been included. To convert to the time domain from the frequency domain, the toolbox uses the inverse fast Fourier transform and to convert to the frequency domain from the time domain, the toolbox uses the fast Fourier transform.

#### 2.4.3 Interleaving Functions

Depending on the artifact and the application, it may be desirable to insert an artifact across edit-ON and edit-OFF transients, for example to mimic frequency drift across an acquisition. Alternatively, it may be desirable to restrict an artifact to specific subspectral data. The *interleave()* function takes in the edit-ON and edit-OFF transients as separate matrices and interleaves them into a single matrix starting with edit-ON while the *undo_interleave()* function takes in the single interleaved matrix and returns the edit-ON and edit-OFF transient matrices.

#### 2.4.4 Scaling Functions

To facilitate the use of different data that will have been generated with different scales, data can be scaled prior to applying the artifact functions. A pair of functions to scale and rescale the provided spectral data has been included. The scaling function normalized the individual frequency domain transients by the maximum absolute value of the complex spectrum. This second function returns the data back to its original scale after artifacts have been applied.

## 3. RESULTS

In Figures 2 – 8, each of the simulated artifacts are shown on the difference spectrum, which has been generated from a single ON-OFF pair with varying degrees of severity while Figure 9 shows a linear frequency drift across multiple ON-OFF pairs. The artifact-free spectrum is also shown in Figures 2-9. Figure 9 shows a linear frequency drift across multiple ON-OFF pairs along with the artifact-free spectrum.

**Figure 2.**
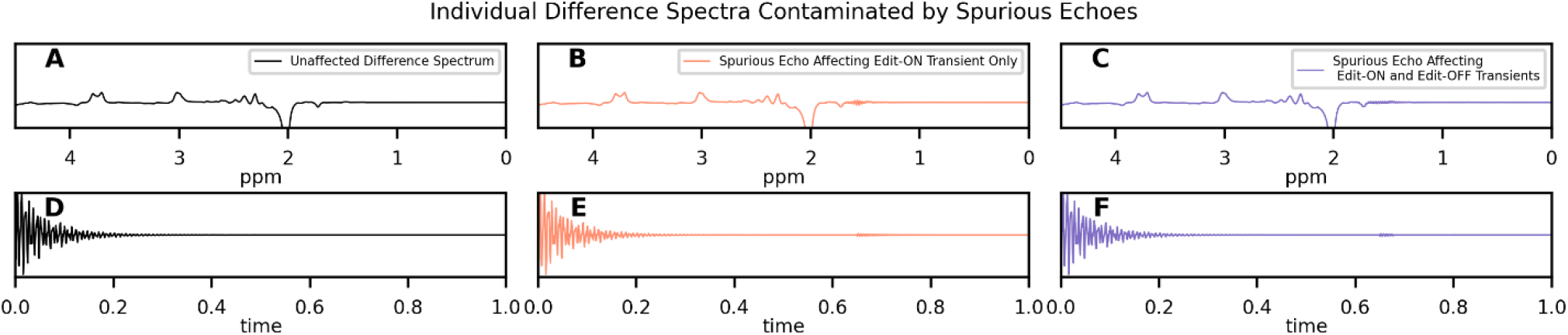
Example spurious echo artifacts generated from the *add_spur_echo_artifact()* function. Frequency domain (top, **A-C**) and time domain (bottom, **D-F**) representations are shown. (**A** and **D** black) Difference spectrum with no spurious echo artifact. (**B** and **E** orange) Difference spectrum with spurious echo artifact with a chemical shift of 1.5 ppm applied to the edit-ON transient. (**C** and **F**, purple) Difference spectrum with spurious echo artifact applied to both the edit-ON and edit-OFF transient pair, also to appear at 1.5 ppm and a ±5% difference in parameter values for the edit-OFF transient.

**Figure 3.**
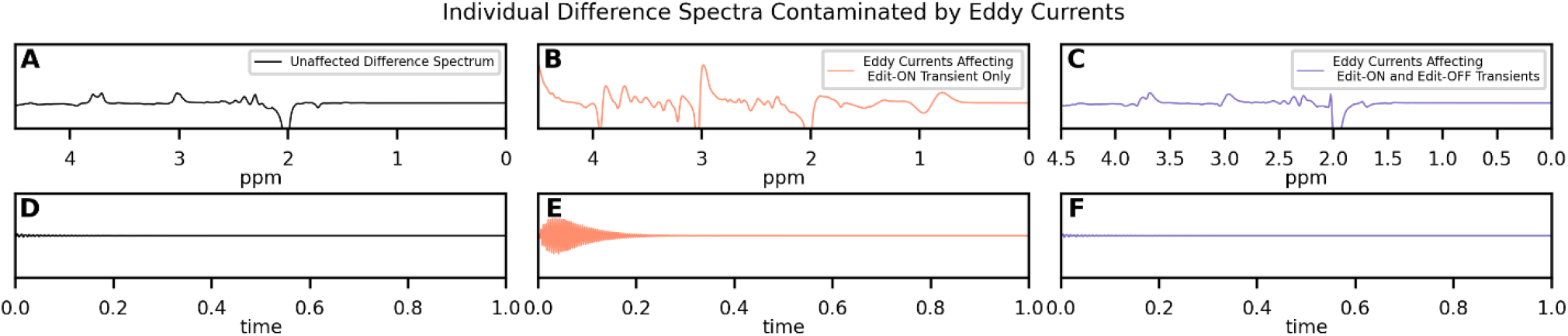
Example of an eddy current artifact, generated with the *add_eddy_current_artifact()*. Frequency domain (top, **A-C**) and time domain (bottom, **D-F**) representations shown. (**A** and **D**, black) Difference spectrum with no artifact added. (**B** and **E** orange) Difference spectrum with eddy current artifact applied to the edit-ON transient. (**C** and **E** purple) Difference spectrum with eddy current artifact applied to both the edit-ON and edit-OFF transient pair. The same amplitude and time constants were used for both edit-ON artifacts (**B**/**E** and **C**/**F**) and ± 0.5% difference in parameter values for the edit-OFF transient in **C**/**F**.

**Figure 4.**
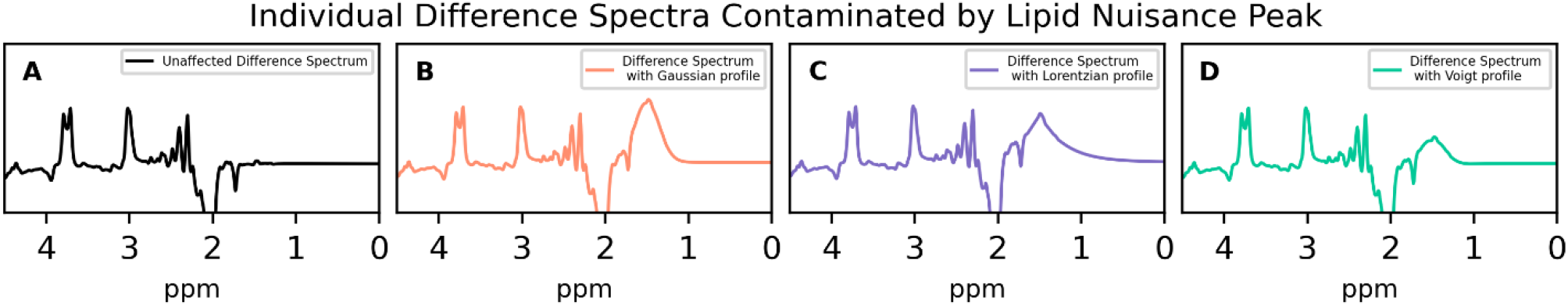
Example of a nuisance peak at 1.5 ppm being simulated using *add_nuisance_peak()*. (**A** black) Difference spectrum with no added lipid peak. (**B** orange) Difference spectrum with Gaussian shape profile selected for the nuisance peak. (**C** purple) Difference spectrum with Lorentzian shape profile selected for the nuisance peak. (**D** green) Difference spectrum with pseudo-Voight shape profile selected for the nuisance peak. For all nuisance peaks, the peak was centered at 1.5 ppm with the tails affecting the surrounding baseline.

**Figure 5.**
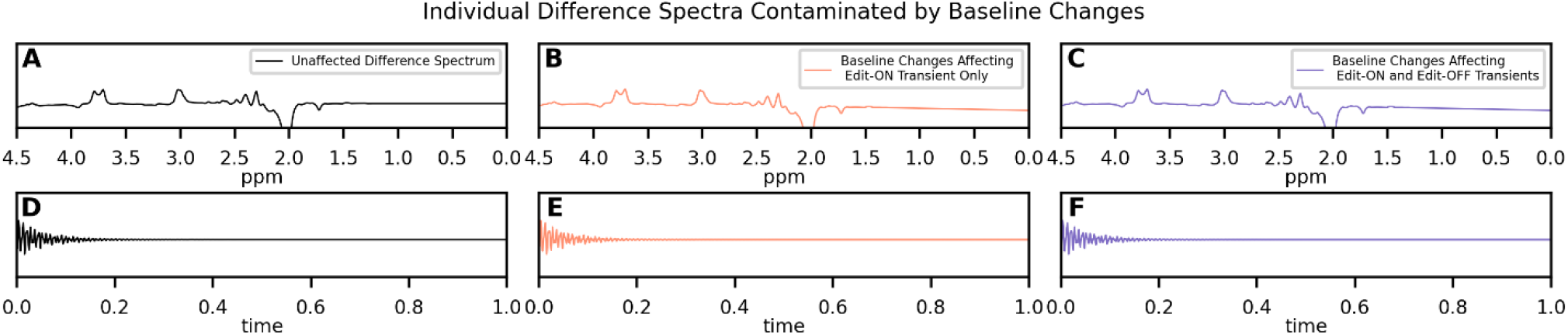
Example baseline contamination simulation using *add_baseline()*. Frequency domain (top, **A-C**) and time domain (bottom, **D-F**) representations shown. (**A** and **D** black) Difference spectrum with no baseline contamination. (**B** and **E**, orange) Difference spectrum with three summed sine wave functions added to the edit-ON transient only. (**C** and **F** purple) Difference spectrum with the same three summed sine wave bases added to both the edit-ON and edit-OFF transient pair and a ±0.5% difference in parameter values for the edit-OFF transient.

**Figure 6.**
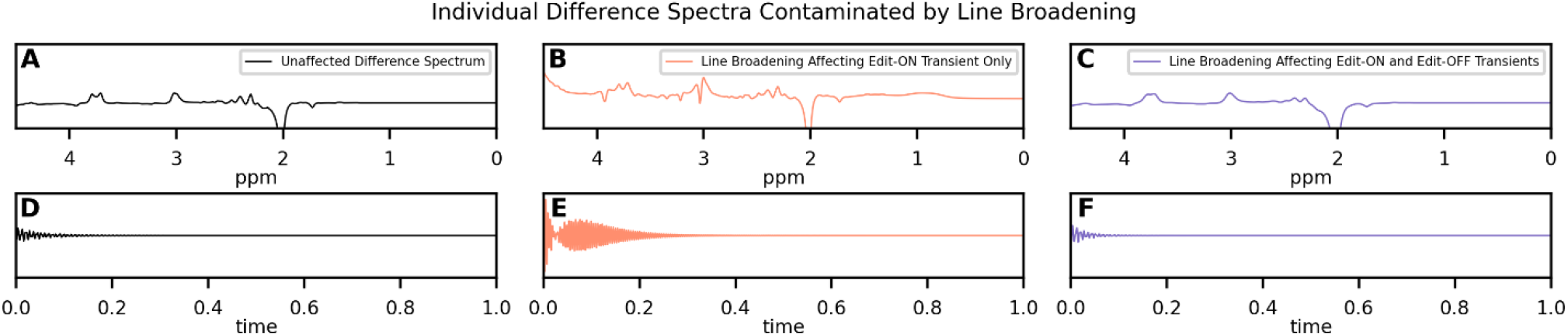
Example of line broadening using of *add_linebroad()* function. Frequency domain (top, **A-C**) and time domain (bottom, **D-F**) representations shown. (**A** and **D** black) Difference spectrum with no line broadening. (**B** and **E** orange) Difference spectrum with line broadening added to the edit-ON transient only. (**C** and **F** purple) Difference spectrum with the same parameter values as in (**B/E**, orange) added to both the edit-ON and edit-OFF transient pair and a ±0.5% difference in parameter values for the edit-OFF transient.

**Figure 7.**
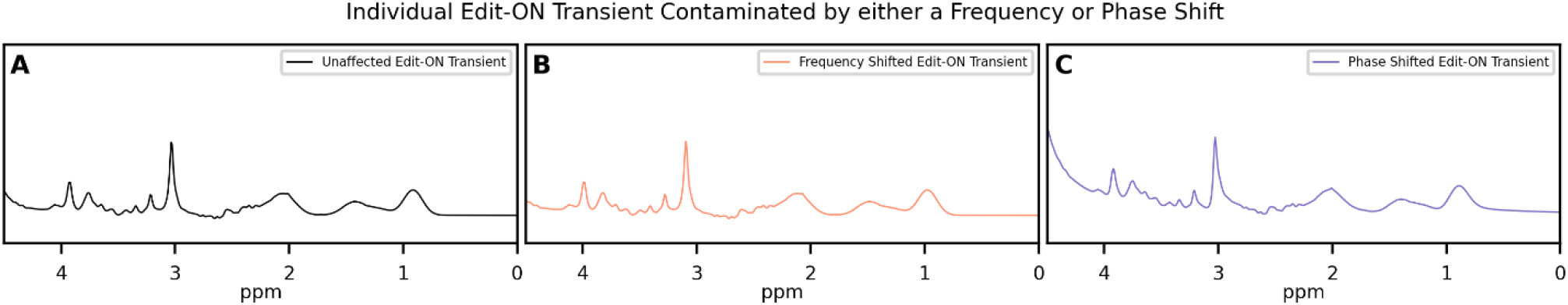
Example of simulated sporadic motion from frequency and phase shifts using *add_freq_shift()* and *add_zero_order_phase_shift()* functions. (**A** black) Single edit-ON transient not affected by frequency or phase shifts. (**B** orange) Single edit-ON transient affected by a large frequency shift (20 Hz). (**C** purple) Single edit-ON transient affected by a large phase shift (60 degrees).

**Figure 8.**
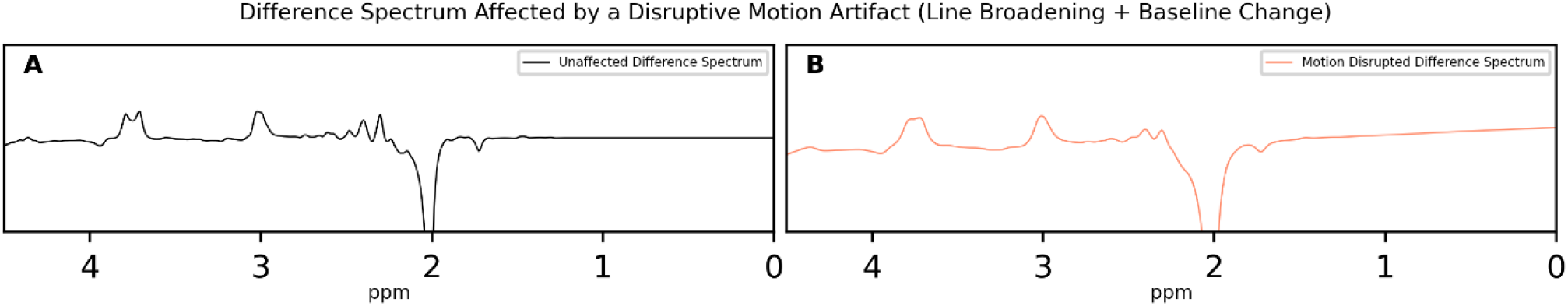
Example of simulated disruptive motion artifact generated using *add_disruptive_motion_artifact()* which uses *add_linebroad()* and *add_baseline()* functions. (**A** black) Difference spectrum with no disruptive motion artifact applied. (**B** orange) Difference spectrum with disruptive motion artifact applied with default amplitude and variance parameter for *add_linebroad()* and default motion profile for *add_baseline()* function.

**Figure 9.**
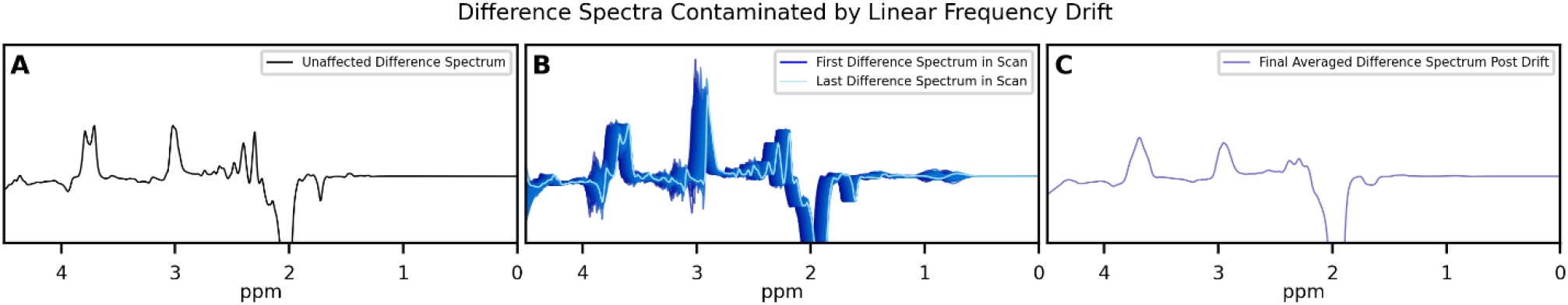
Example of linear drift to mimic progressive head motion or gradient induced frequency drift using the *add_freq_drift_linear()* function. (**A** black) Difference spectrum without any frequency drift. (**B** blue) Difference spectra with a 15 Hz linear frequency drift across 318 transients. The first difference transient is in dark blue and the last difference transient affected by the drift is in light blue. Intermediate difference transients form a gradient from dark blue to light blue. (**C** purple) The final averaged difference spectrum.

There are use cases in which the user should be cautious about the order of adding multiple artifacts and the magnitude of parameter values. For example, as seen in Figure 10a, the order of operations can impact spurious echoes. When adding complex noise and spurious echoes, if the complex noise is added first, the spurious echo will have a greater amplitude than if complex noise is added after the spurious echo.

**Figure 10.**
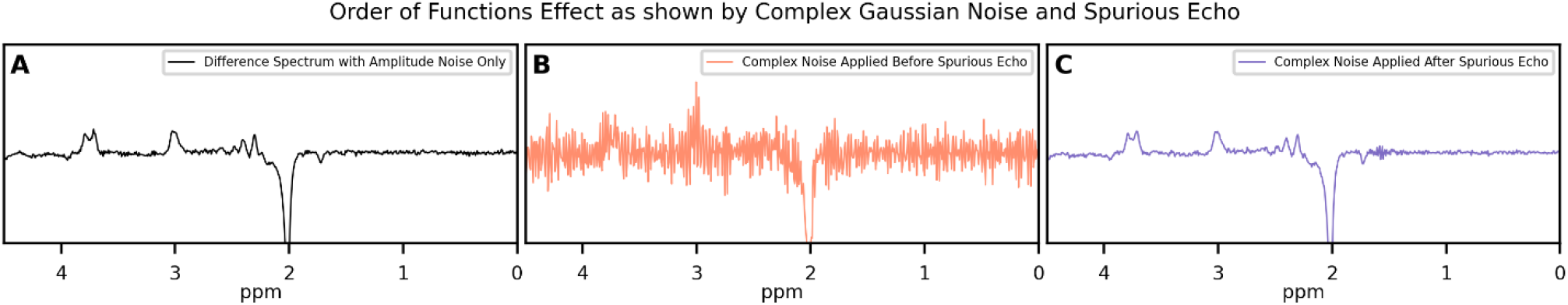
Considerations when using the toolbox. (**A-C**) Example to illustrate the impact of the order of applied functions as seen with *add_time_domain_noise()* and *add_spur_echo_artifact()*. (**A** black) Difference transient with only *add_time_domain_noise()*. (**B** orange) Difference spectrum where complex Gaussian noise was added prior to a spurious echo artifact resulting in significant visual knots or nulled regions, particularly apparent between 0 and 2 ppm. (**C** purple) Difference spectrum where complex Gaussian noise was added after the spurious echo without the same degree of knots.

## 4. DISCUSSION

Simulations of GABA+-edited MRS data have previously been important to understanding spectral behavior (e.g., ref [26]). Simulated data has shown utility for developing preprocessing methods such as frequency and phase correction [27-29]. Furthermore, with the increase in machine learning being applied in spectroscopy [13], comprehensive tools to augment simulated data by incorporating simulated artifacts to better represent *in vivo* data are required. However, to our knowledge, there is not a comprehensive toolbox that includes multiple artifacts commonly encountered in GABA+-edited MRS. The *SMART MRS* simulated artifact toolbox works to fill this gap by providing a set of Python functions that can easily integrate with the user’s existing framework for the simulation of commonly occurring artifacts with flexible user parameters. In addition, artifacts to simulate were selected based on occurrence, difficulty to remove through preprocessing, and probability of causing a transient or entire scan to be discarded [7-10,14]. While it was originally developed for simulated GABA+-edited MRS data, it is adaptable and can be modified to simulate other spectroscopy data, including non-edited MRS or macromolecule-suppressed GABA. In addition, given the predominance of Python in machine learning applications, and the flexibility of Python, we expect this will accommodate most applications.

There are, however, comparisons to be made with existing software. While FID-A [4] is an extensive package with tools for RF pulse simulation, spectrum simulation and spectrum processing, it is MATLAB-based making it more computationally expensive to interface with when exclusively dealing with Python code. Moreover, it does not provide as many artifacts as *SMART MRS* (such as spurious echoes or baseline contamination). VESPA [30] is also an extensive package with similar functionality as FID-A [4] and allows for the simulation of some artifacts but not all, for example there isn’t a function to add eddy currents or spurious echoes. MRS-SIM [20] has the ability to simulate some artifacts, but it does not focus on editing and does not provide functions which seek to combine artifacts into a more complex event such as motion.

The toolbox differentiates itself not only by the variety of artifacts offered but also by its flexible design, which includes setting default parameter values which the user can choose to override or adjust. Additional support functions have also been added to accommodate different inputs. *Interleave()* and *undo_interleave()* functions were created to store data in a sequential way representing an *in vivo* scan where edit-ON and edit-OFF transients are acquired in an interleaved fashion. In addition, the *scale()* function was created to scale complex MRS data so that artifact function default values are within an appropriate range by visually assessing spectrum quality. This is to allow for more realistic artifacts within the data and obviates the need to determine arbitrary parameter values for the user’s unique set of data. If the user wishes to undo the scaling of their data, the *undo_scale()* function simply applies an inverse scaling factor to return the data to its original scale. The toolbox was also designed to accommodate the unique demands of the user allowing them to set the precise number, location and clustering of artifacts.

While the toolbox will enhance the diversity of simulated MRS data by facilitating the integration of some common artifacts, the authors acknowledge some limitations. First, since studies typically present preprocessed (artifact corrected data) and exclude poor, unrecoverable data without further elaborating on the artifacts themselves, the technical descriptions of artifacts are limited. Many artifacts are readily identified on *in vivo* spectra, but mathematics descriptions and realistic parameter values of artifacts are more limited. In this toolbox, the artifact functions were designed to balance the accuracy of replicating the phenomena which led to the artifact and the ease of user-implementation. As such, some of the artifact functions have a reduced number or combined set of features and some of the default parameter values were selected in the context of the definition of ‘poor quality’ while still allowing the user to set their own parameter values. In addition, with limited mathematical resources, the toolbox was validated through visual assessment of artifacts under different sets of conditions (parameter values) and supported by additional sources when possible. Since this work focused on visual validation of 3T single voxel GABA-edited MRS, further validation may be needed for data which falls outside these parameters. This also extends to the order in which artifacts are applied. While practically users can simulate and add artifacts in any order, *in vivo*, artifacts occur concurrently and thus the impact of sequentially adding simulated artifacts is not well known. Thus, artifacts may have additive effects, and the ordering of multiple artifacts should be considered to prevent unintended changes to previously added artifacts (as seen in Figure 10). Depending on the application, users may need to create and compare different pipelines to fully determine the impact of order of the included artifacts.

## 5. CONCLUSION

*SMART MRS* is an open-source Python simulated artifact toolbox for GABA+-edited MRS data which allows users to add spurious echoes, eddy currents, line broadening, baseline changes, linear frequency drift, frequency and phase shifts, nuisance peaks such as lipid contamination and different degrees of motion to simulated MRS data. The toolbox is available as a pip installable Python package: SMART_MRS and a set of examples on how to use the toolbox are available at https://github.com/HarrisBrainLab/SMART_MRS. The *SMART MRS* artifact toolbox was designed to be an open-source tool for users to improve upon as research continues in the field. Suggestions and contributions can be made at the above GitHub link.

## DATA AND CODE AVAILABILITY

This toolbox was developed using Python version 3.11 (with NumPy (v.1.25.2), SciPy (v.1.11.4), and Nibabel (v.5.2.1) dependencies), and MATLAB version 2021b and FID-A (v.1.2). The toolbox is available as pip installable SMART_MRS and examples of how to use the toolbox can be found at the following GitHub repository: https://github.com/HarrisBrainLab/SMART_MRS.

## ACKNOWLEDGEMENTS

This work was presented in part by Bugler et al. as “*Artifact Simulation Toolbox for GABA-Edited Magnetic Resonance Spectroscop*y” at the International Society for Magnetic Resonance in Medicine Annual Meeting 2024 in Singapore.

